# Dynamically regulated Focal adhesions coordinate endothelial cell remodelling in developing vasculature

**DOI:** 10.1101/2022.02.01.478747

**Authors:** Tevin CY. Chau, Teodor E. Yordanov, Jason A. da Silva, Scott Paterson, Alpha S. Yap, Benjamin M. Hogan, Anne Karine Lagendijk

**Author notes:** Correspondence should be addressed to AKL.

## Abstract

The assembly of a mature vascular network involves the coordinated control of cell shape changes to regulate morphogenesis of a complex vascular network. Cellular changes include a process of endothelial cell (EC) elongation which is essential for establishing appropriately sized lumens during vessel maturation^1-3^. However how EC elongation is dynamically regulated *in vivo* is not fully understood since live monitoring of this event can be challenging in animal models. Here, we utilise the live imaging capacity of the zebrafish to explore how integrin adhesion complexes, known as Focal Adhesions (FAs), control EC dynamics in live flow pressured vasculature. To do this, we generated a zebrafish mutant, deficient for the integrin adaptor protein Talin1. Notably, unlike the severe cardiovascular defects that arise in Talin1 knockout mice^4^, vasculogenesis still occurs normally *talin1* mutants and cardiac output remains sufficient up to two days post fertilisation (dpf). This allowed us to uncouple primary roles for FAs in ECs during subsequent morphogenesis events, including angiogenesis and vessel remodelling, without interference of secondary effects that might occur due to systemic vessel failure or loss of blood flow. We further established a FA marker line, expressing endothelial Vinculin-eGFP, and demonstrated that FAs are lost in our *talin1* mutants. This Vinculin transgene represents the first *in vivo* model to monitor endothelial FA dynamics. Loss of FAs in *talin1* mutants, leads to compromised F-actin rearrangements, which perturb EC elongation and cell-cell junction linearisation during vessel remodelling. Chemical induction of actin polymerisation can restore these cellular phenotypes, suggesting a recovery of actin rearrangements that are sufficient to allow cell and junction shape changes. Together, we have identified that FAs are essential for active guidance of EC elongation and junction linearisation in flow pressured vessels. These observations can explain the severely compromised vessel beds, haemorrhage and vascular leakage that has been observed in mouse models that lack integrin signalling^4-8^.

## INTRODUCTION

Cells are mechanically linked to the extracellular matrix (ECM) via dynamic protein complexes, whereby integrin heterodimers form the main ECM ligand binding receptors that couple adhesion to the actin cytoskeleton^9,10^. Integrins, however, cannot bind to actin directly, and full mechanical linkage is achieved via adaptor proteins such as talins^9-13^. Talins are recruited to ligand-bound integrins and mechanical tension applied externally or from the actin cytoskeleton can unfold talin to reveal additional binding sites for other proteins, such as vinculin^12^. Via this direct and indirect recruitment of numerous cytosolic proteins, cell-ECM adhesion structures are formed that control both biochemical and biophysical signalling. Further clustering of integrins, actin bundling, and re-enforcement of these interactions lead to the maturation of cell-ECM adhesion sites, referred to as Focal Adhesions (FAs)^14^. Talin is required for FA formation, actin-attachment, and FA maturation^12,15^.

Since the initial discoveries of cardiovascular defects in mouse models with compromised integrin signalling^16^, vascular biologists have had a great interest in understanding how FAs alter vessel growth, function and stability. At the receptor level, the integrin-β1 subunit, as well as the α5/αv subunits, have been proven to be essential for angiogenesis and vascular integrity^17-20^. Underpinning the relevance of talins for integrin function, endothelial specific knock-out (KO) of Talin1 (the main Talin protein expressed by ECs^4,21^) results in similar phenotypes^4,22^, including reduced vessel network complexity and haemorrhaging. Likewise, Talin1-deficient zebrafish develop haemorrhages in the brain, however vascular morphogenesis has not been analysed in detail in this zebrafish model^23^. It is important to note that FAs do not exist and function as isolated adhesive structures, instead significant crosstalk occurs with cadherin-based adhesions at cell-cell junctions^24-26^. Crosstalk happens by reciprocal binding to the actin cytoskeleton, and via recruitment of identical intracellular effector proteins. Haemorrhaging of FA deficient vasculature therefore has been suggested to be a consequence of impaired crosstalk, since disrupted VE-cadherin localisation has been identified in retinal vasculature of integrin-β1 and Talin1 KO mice^22,27^. However, the mechanisms underlying this crosstalk remains to be determined.

Together, landmark studies using mouse KOs have clearly shown that FAs are essential for vascular morphogenesis and vessel function. However, phenotypic analyses of early vascular morphogenesis were performed in constitutive EC KOs which develop severe cardiac and vascular abnormalities that lead to embryonic lethality^4-8^. It is well appreciated that FA assembly and function is also dependent on cell extrinsic factors including flow shear stress^28-31^. The severity of the mouse KO phenotypes might therefore have complicated the identification of primary cellular processes that require FA function during these early stages of vascular network formation. In this study, we sought to better understand the early vascular impact of FAs, using a novel zebrafish *talin1* mutant model, *tln1*^*uq1al*^. In *tln1*^*uq1al-/-*^mutants cardiac output remains stable, and thus blood circulation persists, until two days post fertilisation (2 dpf) providing a window of opportunity to study early vascular events when blood flow is preserved. In other words, this model provides a unique opportunity to study the EC-specific consequences of compromised FA function in flow pressured vasculature of live animals.

First, we utilised a novel FA marker line, expressing a vascular restricted Vinculin-eGFP fusion protein, to show that *tln1*^*uq1al-/-*^mutants lack mature, Vinculin-positive, FAs. Further, high-resolution live-imaging of VE-cadherin and F-actin uncovered that FAs are required for EC elongation and cell-cell junction linearisation during angiogenesis and vascular remodelling. Mechanistically, enhancing actin polymerisation restores EC elongation and junction linearisation of FA deficient ECs. Together, this work has identified that in ECs that experience normal blood flow, FAs are required for vessel remodelling by driving actin rearrangements that are translated into EC elongation and linearisation of EC junctions. These defects in EC elongation and junction linearisation likely contribute to the severely compromised vessel beds, haemorrhage and vascular leakage that has been observed in mouse models that lack integrin signalling.

## RESULTS

### Establishing in vivo models to study focal adhesion function and dynamics during blood vessel morphogenesis

To examine the role of cell-ECM coupling in vascular morphogenesis, we generated a *talin1* mutant line, *tln1*^*uq1al*^. We designed a guide RNA that would target the highly conserved Talin1 F1 FERM-subdomain (Sup Fig. 1A). This domain is known to be essential for integrin binding^11,12^. The *tln1*^*uq1al*^ allele harbours an in-frame deletion of 6 amino acids and substitution of one amino acid (Fig. 1A, B). The mutation is located in the F1 FERM domain and, based on the predicted structure and location of the missing amino acids using Alphafold2^32,33^, we hypothesize that our mutant Talin1 protein is highly likely to alter protein folding, which would prevent integrin binding and thus FA formation (Sup Fig. 1B). We further did not detect evidence for genetic compensation^34^ by other Talins at the transcriptional level (Sup Fig. 1C). Phenotypically, the first time-point at which we could distinguish *tln1*^*uq1al-/-*^mutants from siblings was at two days post fertilisation (2 dpf). The mutants present with cranial haemorrhaging and a mild cardiac edema (Fig. 1C), comparable to haemorrhaging phenotypes in other mice and zebrafish knockout models for FA components^4-8,22,35-38^.

**Figure 1:**
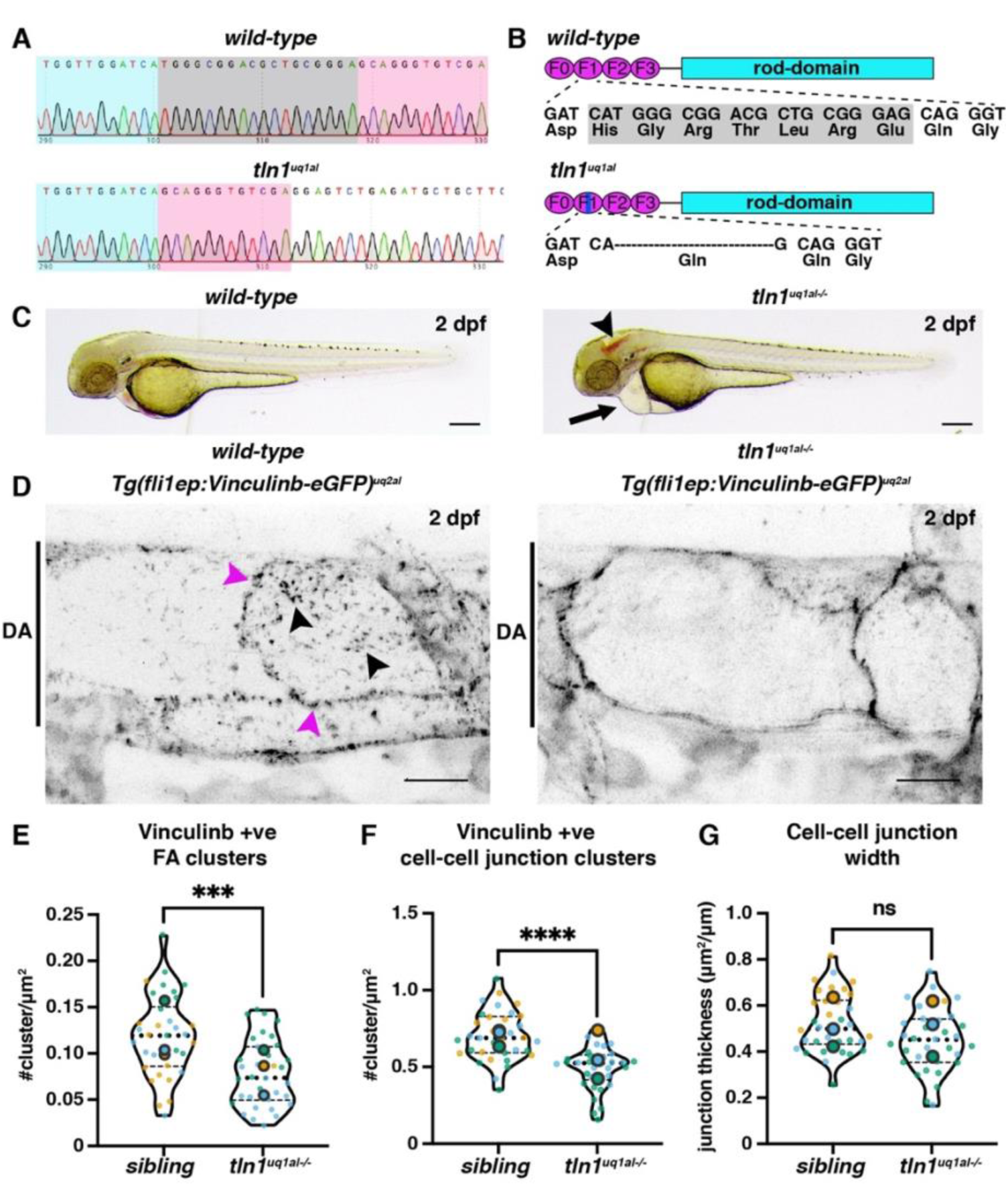
Novel zebrafish models to study focal adhesion function and dynamics in live vasculature. (A) Chromatograms of genomic sequencing of a *wild-type* sibling embryo (top) and a homozygous *tln1*^*uq1al-/-*^mutant (bottom). Bases highlighted in grey in the wild-type are deleted in *tln1*^*uq1al-/-*^mutants (B) Schematics of Talin1 protein showing with the 18bp deletion in *tln1*^*uq1al-/-*^resulting in an in-frame deletion of 6 amino acids (His-Gly-Arg-Thr-Leu-Arg) and a Glu to Gln substitution in the F1 FERM domain. (C) Brightfield images showing body morphology of a *wild-type* sibling versus a *tln1*^*uq1al-/-*^mutant at 2 dpf. Arrowhead indicated cranial haemorrhaging in the mutant. Arrow indicates cardiac edema in the mutant. Scale bar = 100µm (D) Expression of Vinculinb at FAs (magenta arrowheads) and at cell-cell junctions (black arrowheads) in the dorsal aorta (DA) of a *wild-type* sibling at 2 dpf (left). In *tln1*^*uq1al-/-*^mutants (right), expression at FAs in greatly reduced, whilst a diffuse presence of Vinculinb is observed at cell-cell junctions. Scale bar = 10µm (E) Number of Vinculinb-eGFP positive FA clusters. (F) Number of Vinculinb-eGFP positive clusters at cell-cell junctions. (G) Quantification of cell-cell junction width based on Vinculin-eGFP expression. All graphs present n=3 biological replicates, n=34 siblings and n=34 *tln1*^*uq1al-/-*^mutants. Replicate averages are depicted by large circles and smaller circles present individual data points of each replicate (colour matched).

Talin1 is essential for Vinculin recruitment and formation of mature FAs^11,12^. To explore Vinculin recruitment and formation of FAs in our mutant zebrafish, we generated a novel vascular restricted Vinculin transgenic line. Zebrafish have two vinculin paralogs, *vinculina* and *vinculinb*, that are functionally redundant and share equal sequence conservation with mammalian Vinculin^39^. We tagged Vinculinb with eGFP at the C-terminus and generated a vascular restricted transgenic line to visualise Vinculin dynamics, *Tg(fli1ep:Vinculinb-eGFP)*^*uq2al*^. Live imaging of the main axial artery, the dorsal aorta (DA), in wild-type zebrafish embryos, revealed Vinculinb localization in distinctive and highly dynamic clusters at cell-cell junctions and at FAs (Fig. 1D and Movie 1). To our knowledge, this represents the first *in vivo* model that reports FA dynamics in ECs of live vasculature. Vinculinb localization at FAs was strongly reduced in *tln1*^*uq1al-/-*^mutant embryos (Fig. 1D-E and Movie 1). Notably, Vinculinb could still be detected at *tln1*^*uq1al-/-*^mutant cell-cell junctions. However, whilst in wild-type embryos we uncovered a distinctive punctate pattern, in *tln1*^*uq1al-/-*^mutants Vinculinb redistributed more continuously along the junctions, with no significant difference in the overall width of the junctions (Fig. 1D-G and Movie 1). This data not only supports our hypothesis that our *tln1*^*uq1al-/-*^allele is a null, it also verifies that mature Vinculin-positive FAs fail to form in the absence of Talin1. Our zebrafish model therefore is uniquely suited to explore the consequences of FA loss in live flow pressured vasculature.

### Talin1 function is required for efficient sprouting angiogenesis

F-actin-rich filopodial extensions have been shown to be essential to facilitate efficient sprouting angiogenesis of intersegmental vessel (ISVs) in zebrafish^1^ and anchorage of such filopodia to the ECM occurs via FAs. We therefore sought to evaluate angiogenic sprouting efficacy in our *tln1*^*uq1al-/-*^mutants. To label the vasculature, we made use of our previously reported VE-cadherin transgenic line^*40*^, *Tg(ve-cad:ve-cadTS)*^*uq11bh*^ (further referred to as VE-cadherin-TS). Live imaging of the zebrafish trunk revealed a reduction in ISV sprouting in *tln1*^*uq1al-/-*^mutants (Fig. 2A-B). To determine if altered filopodia dynamics was responsible for this, we examined filopodia number and length, using a transgenic line that expresses a membranous mCherry; *Tg(kdrl:Hsa*.*HRAS-mCherry)*^*s916*^. Both the number and length of ISV filopodia at the sprouting tip were comparable between *tln1*^*uq1al-/-*^mutants and siblings (Fig. 2C-E), suggesting that other cellular processes, such as EC elongation, might be responsible for the mild reduction in sprout length.

**Figure 2:**
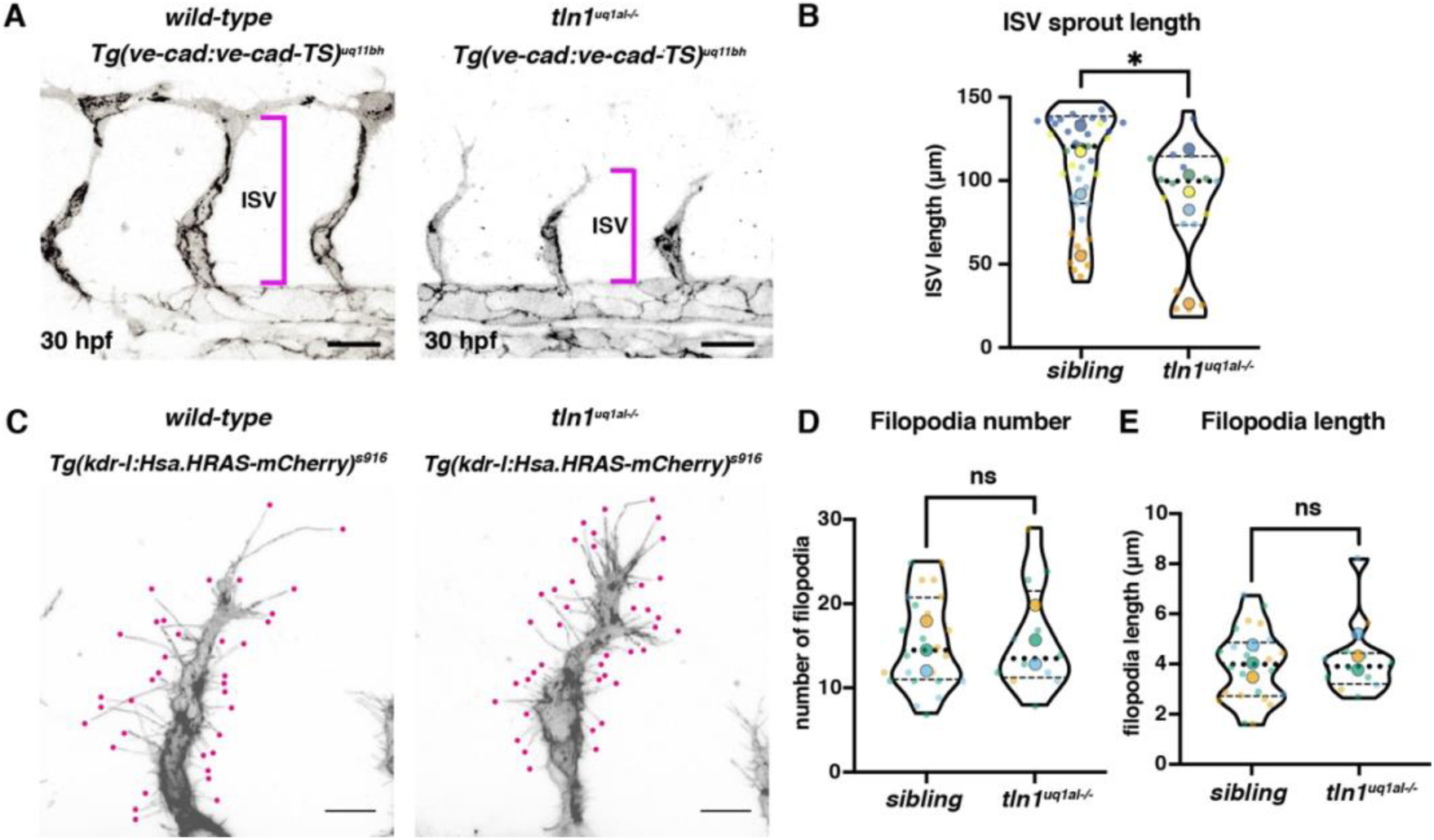
Focal adhesion function is required for efficient angiogenic sprouting. (A) Trunk vasculature of *Tg(ve-cad:ve-cad-TS)* positive wild-type sibling and in *tln1*^*uq1al-/-*^mutant embryo at 30 hpf. ISV sprouts are indicated by magenta brackets. Scale bar = 50µm (B) Quantification of ISV sprout length, showing a mild reduction in mutants, 5 biological replicates, n=30 siblings and n=19 *tln1*^*uq1al-/-*^mutants. (C) High resolution live image showing filopodial extensions at the sprouting front of an ISV in a wild-type sibling and in *tln1*^*uq1al-/-*^mutant embryo at 30 hpf. Embryos express *Tg(kdrl:Hsa*.*HRAS-mCherry)*^*s916*^, labelling the EC membrane (magenta). Scale bar = 10µm (D,E) Quantification of filopodia number (D) and length (E) at the tip of ISV sprouts, n=3 biological replicates, n=24 siblings and n=12 *tln1*^*uq1al-/-*^mutants. In all graphs replicate averages are depicted by large circles. Smaller circles present individual data points of each replicate (colour matched).

### Blood vessel remodelling involves focal adhesion guided elongation of endothelial cells

Upon sprouting angiogenesis, the initial framework of the trunk vasculature is established, and the vasculature subsequently undergoes a process of EC remodelling^41^. EC elongation is one of the main remodelling events that contributes to maturation of embryonic vasculature^40,42-45^.

To investigate EC remodelling, we again made use of our VE-cadherin-TS transgenic line and performed live imaging at 50 hpf. Importantly, cardiac function was still sufficient at this stage in *tln1*^*uq1al-/-*^mutants, and thus the ECs are experiencing blood flow (Movie 2). Initial analysis of the vascular network at 50 hpf revealed that, despite reduced angiogenesis efficiency, the ISVs in *tln1*^*uq1al-/-*^mutants do reach the dorsal side of the trunk and establish the dorsal longitudinal anastomotic vessel (DLAV) (Fig. 3A). However, the dorsally derived DLAV plexus fails to fully form (Fig. 3A, B). We further identified that *tln1*^*uq1al-/-*^mutant ISVs frequently regressed and disconnected from the main vessel network (Fig. 3A, C and Movie 3), suggesting that the network is not maintained. Based on this analysis, we hypothesized that although the primitive framework of DA, posterior cardinal vein (PCV), ISVs and DLAV is established, the network is not maintained.

**Figure 3:**
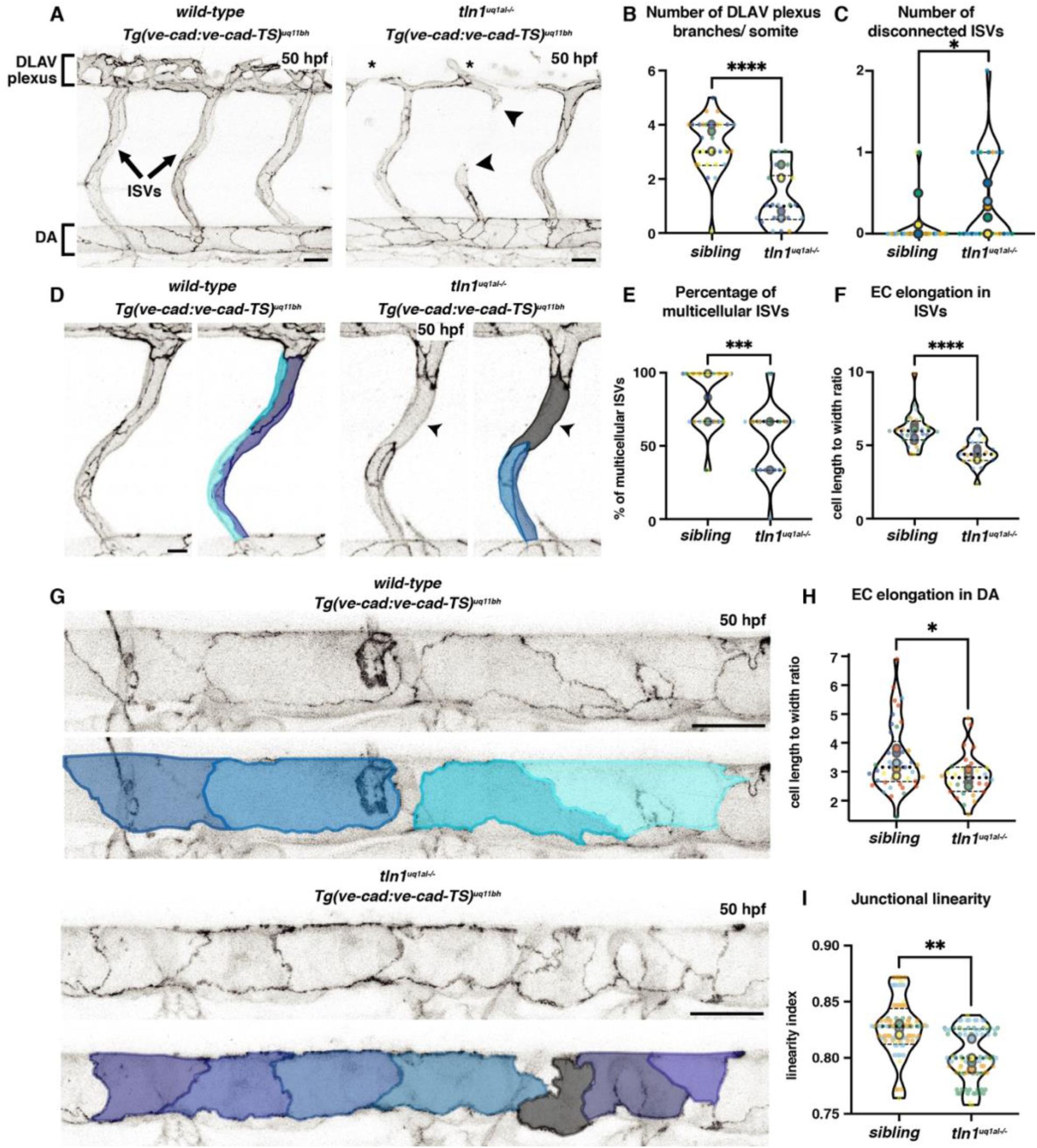
Loss of Focal Adhesions results impairs endothelial cell remodelling and vessel network maintenance. (A) Trunk vasculature of *Tg(ve-cad:ve-cad-TS)* positive wild-type sibling and in *tln1*^*uq1al-/-*^mutant embryo at 50 hpf, showing a reduced of DLAV plexus formation (asterisk) and disconnected ISVs (arrowheads) in the mutant. Scale bar = 50µm (B) Quantification of the number of DLAV branches per somite, n=5 biological replicates, n=29 siblings and n=22 *tln1*^*uq1al-/-*^mutants. (C) Quantification of the number of disconnected ISVs across the width of two somites, located above the yolk extension, n=5 biological replicates, n=29 siblings and n=22 *tln1*^*uq1al-/-*^mutants. (D) Left: Single ISVs of a wild-type sibling and in *tln1*^*uq1al-/-*^mutant embryo at 50 hpf, expressing VE-cadherin *Tg(ve-cad:ve-cad-TS)*. Right: Individual ECs false-coloured to indicate EC shapes. Arrowhead indicates a region of the ISV that is uni-cellular in the mutant. Scale bar = 10µm. (E) Quantification of the percentage of multi-cellular ISVs per two somites, n=5 biological replicates, n=31 siblings and n=22 *tln1*^*uq1al-/-*^mutants. (F) Quantification of EC elongation in ISVs, n=5 biological replicates, n=31 siblings and n=22 *tln1*^*uq1al-/-*^mutants. (G) Top: Dorsal aorta (DA) of a wild-type sibling and in *tln1*^*uq1al-/-*^mutant embryo at 50 hpf, expressing VE-cadherin *Tg(ve-cad:ve-cad-TS)*. Bottom: False-coloured duplicate to indicate EC shape. Scale bar = 25µm (H) Quantification of EC elongation in the DA, n=6 biological replicates, n=41 siblings and n=30 *tln1*^*uq1al-/-*^mutants. (I) Quantification of junctional linearity of ECs in the DA, n=6 biological replicates, n=41 siblings and n=30 *tln1*^*uq1al-/-*^mutants. In all graphs replicate averages are depicted by large circles. Smaller circles present individual data points of each replicate (colour matched).

Remodelling of the ISVs and the DA involves a dynamic process of EC elongation^40,42-45^. Of note, EC elongation is involved in the transformation of pre-mature unicellular ISVs into multicellular ISVs^43-45^. In unicellular ISVs, ECs are stacked one on top the other. Upon EC rearrangements, multicellular tubes are formed with multiple cells lining the vessel lumen, which improves vessel stability^43-45^. By utilising the VE-cadherin-TS marker, to outline individual ECs, we found that segments of *tln1*^*uq1al-/-*^ISVs displayed a unicellular configuration at 50 hpf whilst ISVs in siblings progressed had into a multicellular configuration (Fig. 3D, E). Quantification of cell shape revealed that the ECs that made up the *tln1*^*uq1al-/-*^ISVs were less elongated (Fig. 3D, F). Since EC elongation is a major event during multicellular tube formation, we suggest that the reduced capacity of *tln1*^*uq1al-/-*^ECs to elongate results in regions of immature and unstable ISVs, which can explain the ISV regression phenotype (Fig. 3A, C and Movie 3).

Next, we explored whether EC elongation was compromised more broadly in *tln1*^*uq1al-/-*^mutant vasculature by examining EC elongation of cells in the DA. We and others have reported that ECs in the DA undergo a process of EC elongation which is initiated at 2 dpf^40,42^. This process is essential for DA vessel diameter adaptation^42^. Quantification of EC shape in the DA of *tln1*^*uq1al-/-*^and sibling embryos at 50 hpf revealed that *tln1*^*uq1al-/-*^ECs are less elongated (Fig. 3G, H), showing an early defect in DA maturation. An additional hallmark of EC remodelling in the DA is linearisation of EC junctions. In *tln1*^*uq1al-/-*^mutants the ECs in the DA also present with significantly less linear junctions (Fig. 3G, I). This data is in line with previous studies in mice, where irregular VE-cadherin junctions were also identified^8,35,36^.

Together, this data shows that FAs are essential to facilitate EC elongation during vessel remodelling of both larger calibre vessels and capillaries. FA-deficient ECs are further characterized by immature, irregular, cell-cell junctions, which are likely to contribute to increased vessel permeability, a major phenotypic hallmark of FA loss of function mouse models^4,6,7,22,35,36^.

### Focal adhesions facilitate endothelial cell elongation cell-autonomously

We next examined whether the failure of EC elongation was due to a loss of Talin1 function in ECs. We performed cellular transplantation experiments at blastula stages and transferred cells from either *tln1*^*uq1al-/-*^or sibling donor embryos into *Tg(kdrl:Hsa*.*HRAS-mCherry)*^*s916*^ wild-type hosts (Fig. 4A-B). We utilised the VE-cadherin-TS marker, which was only expressed by donor embryos, and measured EC shape within mosaic clones in the DA. We focussed on the DA for these experiments since EC elongation in the DA is prominent from 2 dpf onwards. These quantifications confirmed that ECs require FA function, cell-autonomously, to facilitate their elongation during vessel remodelling (Fig. 4C-E). Notably, *tln1*^*uq1al-/-*^mutant EC junctions were also less linear in mosaic mutant clones (Fig. 4F). Ubiquitous *tln1*^*uq1al-/-*^mutants develop progressive cardiac failure causing a loss of blood circulation at 3 dpf and thus we did not examine EC remodelling in *tln1*^*uq1al-/-*^mutants beyond 2 dpf. Generating chimeric embryos by transplantation, however, allowed us to examine the consequences of FA loss in ECs also at later stages. Whilst EC elongation becomes more pronounced in ECs of wild-type origin between 2 to 3 dpf^40,42^, this progressive elongation did not occur in clones of *tln1*^*uq1al-/-*^mutant. In fact, these FA-deficient ECs at 3 dpf were indistinguishable in shape when compared to EC clones derived from sibling embryos at 2 dpf and junctions remain irregular (Fig. 4D-F). These data show that FAs are essential to facilitate ongoing EC elongation and junction linearisation in the DA during vessel remodelling.

**Figure 4:**
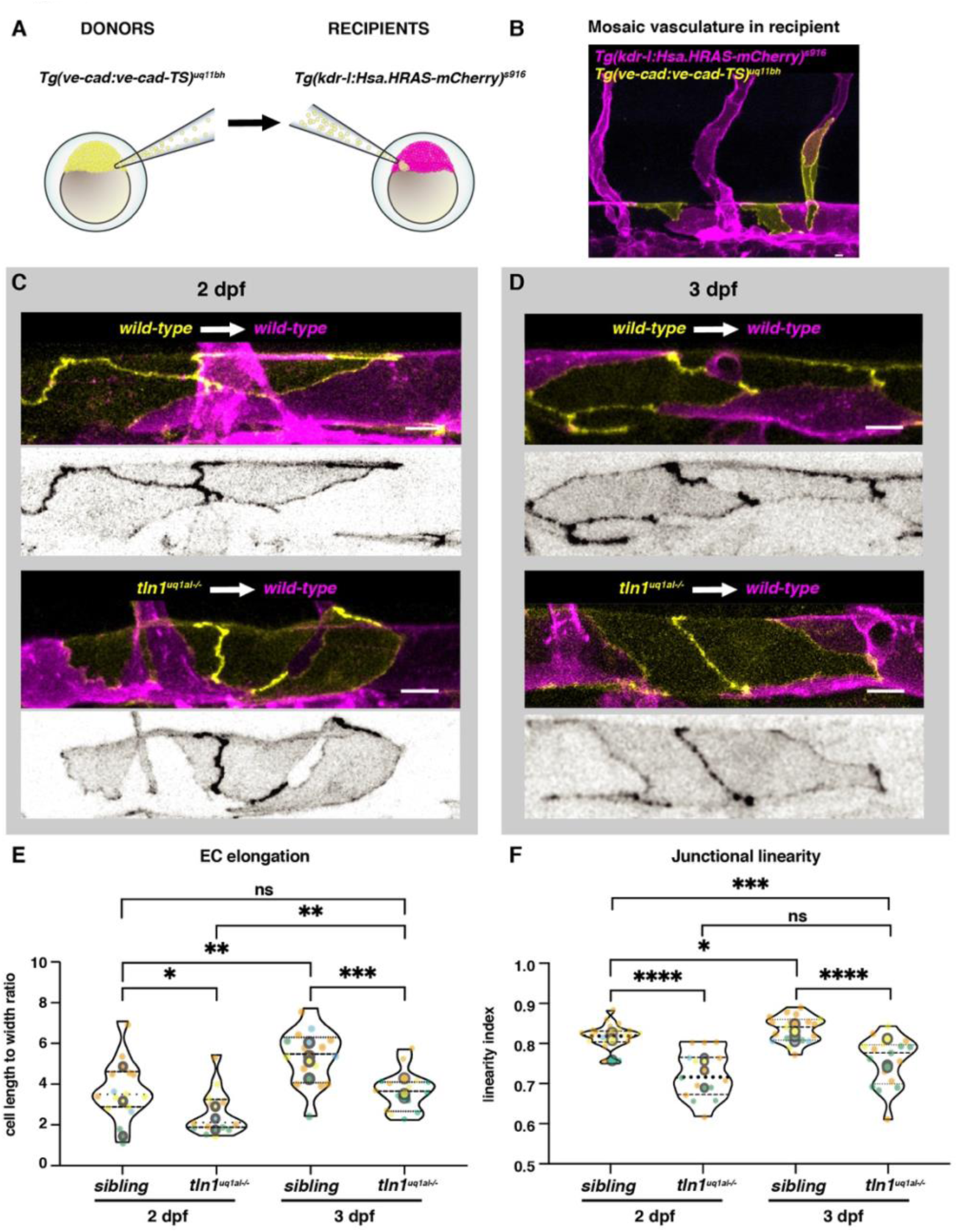
Talin1 is required cell-autonomously to facilitate endothelial cell elongation. (A) Schematic representation of transplantation procedure. For these experiments we transplanted cells at blastula stage from *Tg(ve-cad:ve-cad-TS)*^*uq11bh*^ donors to *Tg(kdrl:Hsa*.*HRAS-mCherry)*^*s916*^ recipients. (B) *Tg(kdrl:Hsa*.*HRAS-mCherry)*^*s916*^ positive (magenta) trunk vasculature of a recipient embryo at 2 dpf with a mosaic clones of cells from the donor, VE-cadherin-TS positive (yellow). Scale bar = 10µm (C) Wild-type dorsal aorta (magenta) of recipient embryos at 2 dpf containing either wild-type donor ECs (top, yellow) or *tln1*^*uq1al-/-*^mutant ECs (bottom, yellow). Scale bar = 10µm. Greyscale images show donor ECs only. (D) Wild-type recipient dorsal aorta (magenta) of recipient embryos at 3 dpf containing wild-type donor ECs (top, yellow) or *tln1*^*uq1al-/-*^mutant ECs (bottom, yellow). Scale bar = 10µm. Greyscale images show donor ECs only. (E) Quantification of EC elongation of transplanted ECs during dorsal aorta maturation from 2 dpf to 3 dpf. (F) Quantification of cell-cell junction linearity of transplanted ECs during dorsal aorta maturation from 2 dpf to 3 dpf. (E,F) n=4 biological replicates, 2 dpf data n=15 siblings and n=14 *tln1*^*uq1al-/-*^mutants, 3 dpf data n=15 siblings and n=16 *tln1*^*uq1al-/-*^mutants. In all graphs replicate averages are depicted by large circles. Smaller circles present individual data points of each replicate (colour matched).

### EC elongation requires forces generated via a functional network of F-actin fibers

Ultimately, dynamic changes in acto-myosin contractility determine cell shape. Vinculin is well known to assist in the reinforcement of FA-actin binding, allowing for enhanced force-bearing. Loss of Vinculin-positive FAs (Fig. 1D, E) thus implies that *tln1*^*uq1al-/-*^mutant ECs are likely to be experiencing a non-physiological distribution of force. We therefore hypothesized that impairment of FAs in *tln1*^*uq1al-/-*^mutants disturbs actin rearrangements, which subsequently hampers the ability of ECs to elongate.

To test this hypothesis, we first examined whether forces generated by polymerized F-actin fibers are required for EC elongation during DA maturation. To do this, we utilised a well-characterised chemical inhibitor of the Arp2/3 complex, CK666^46,47^. We treated wild-type VE-cadherin-TS embryos with CK666 [75μM], from 24 hpf to 50 hpf, and subsequently imaged the DA. Interestingly, CK666 treated embryos presented with bleeding in the brain (Fig. 5A), reduced DLAV plexus (Sup Fig. 2A, B) and multicellular ISV tube formation (Sup Fig. 2A, C). These phenotypes were similar to those we observed in the *tln1*^*uq1al-/-*^mutants (Fig. 1C and Fig. 3A-C). Upon CK666 treatment, ECs in the DA failed to elongate (Fig. 5B-C) and were bounded by immature, irregular, cell-cell junctions (Fig. 5D). The similarity in EC remodelling phenotypes between CK666-treated embryos and *tln1*^*uq1al-/-*^mutants suggested that when FA function is compromised, F-actin movements are perturbed.

**Figure 5:**
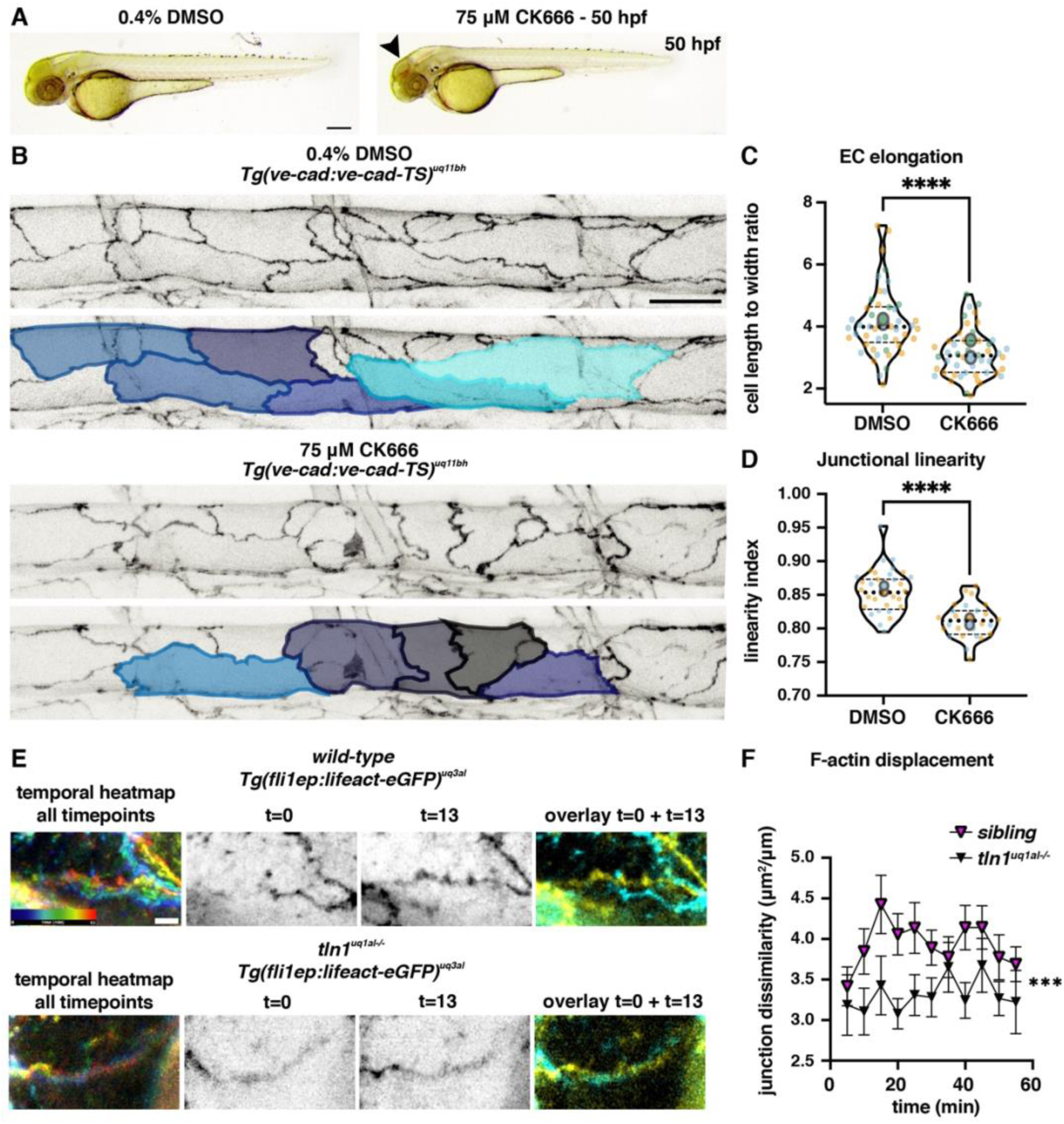
Temporal inhibition of actin polymerisation induces replicated endothelial cell remodelling phenotypes. (A) Brightfield images showing the overall body morphology at 50 hpf of embryos upon a 5-hour treatment with either 0.4% DMSO (control, left) or with CK666 [75uM] (Arp2/3 inhibitor, right). CK666 treated embryos develop cranial haemorrhaging (arrowhead), reminiscent to *tln1*^*uq1al-/-*^mutants (Fig. 1A). Scale bar = 100µm (B) Maximum projection of the dorsal aorta at 50 hpf, directly after treatment with either DMSO (control, top) or CK666 (bottom), showing impaired EC elongation. Duplicate images, one with false-coloured ECs to highlight EC shape. The colours present a spectrum, lighter colours indicate more elongated ECs. Scale bar = 25µm (C) Quantification of EC elongation at 50 hpf, showing a significant reduction upon CK666 treatment, n=3 biological replicates, n=48 DMSO and n=48 CK666 treated embryos. (F) Quantification of cell-cell junction linearity at 50 hpf, showing a significant reduction upon CK666 treatment, n=3 biological replicates, n=48 DMSO and n=48 CK666 treated embryos. (E) Still images form time-lapse imaging (Sup Movie 4). Movie recorded cortical F-actin rearrangements in wild-type (top) and *tln1*^*uq1al-/-*^mutant ECs (bottom) form 48 till 49 hpf. Left: temporal colour-coded projection of 13 timepoints imaged during the hour time-lapse. Middle: Greyscale images of F-actin expression at the start (t=0) and the end of the movie (t=13). Right: overlay of t=0 (cyan) and t=13 (yellow) showing F-actin displacement in the wild-type and very little movement in *tln1*^*uq1al-/-*^mutant. Scale bar = 5μm. (F) Quantification of F-actin displacements of cortical F-actin over the course of a one-hour time-lapse movie in siblings (magenta) and *tln1*^*uq1al-/-*^mutants (black). Each data point represents the average displacement between two consecutive time-points, n=3 biological replicates, n=20 siblings and n=14 *tln1*^*uq1al-/-*^mutants. In C and D replicate averages are depicted by large circles. Smaller circles present individual data points of each replicate (colour matched).

Then, we used a vascular F-actin marker line, *Tg(fli1ep:lifeact-eGFP)*^*uq3al*^, to examine F-actin rearrangements in *tln1*^*uq1al-/-*^animals. Since we have observed changes in cell-cell junction linearisation simultaneous to EC elongation, we hypothesize that FAs drive actin rearrangements that can act on cortical actin fibres near cell-cell junctions. Paatero and colleagues have previously identified junction-based lamellipodia (JBLs) at cell-cell junctions^48^. F-actin protrusions in JBLs bring in junctional proteins, thereby remodelling junction morphology. To generate movies at high temporal resolution, we imaged cortical actin at five-minute intervals for 1 hour starting at 48 hpf (Fig. 5E and Movie 4). In *tln1*^*uq1al-/-*^mutants, the overall displacement of F-actin over the one hour time-lapse was significantly reduced (Fig. 5F), reflecting the failure of cells to elongate. This analysis demonstrated a reduction in F-actin rearrangements in FA-deficient ECs, compared to ECs of sibling embryos. These observations suggest that in absence of FA function, F-actin rearrangements are altered, preventing EC elongation and cell-cell junction elongation.

### Stabilising F-actin can compensate for Focal adhesions loss

We next aimed to determine whether chemical stabilisation of polymerised F-actin in *tln1*^*uq1al-/-*^mutants was sufficient to facilitate EC elongation. To do this, we utilised Jasplakinolide (Jasp), a compound which stabilises actin filaments by reducing disassembly^49^. We treated *tln1*^*uq1al-/-*^mutant and sibling embryos with either Jasp [1μM] or 0.1% DMSO from 24 hpf until 50 hpf, when the embryos were imaged. We observed an improvement in DLAV plexus formation (Sup Fig. 2D, E) and ISVs more frequently matured into multicellular tubes in Jasp treated *tln1*^*uq1al-/-*^mutants (Sup Fig 2D, F), indicating that DLAV sprouting and EC remodelling can be partially restored to FA-deficient ECs by reducing F-actin disassembly.

We further quantified cell shape of ECs in the DA which revealed that EC elongation and junction linearisation was rescued in Jasp treated *tln1*^*uq1al-/-*^mutants (Fig. 6A-C). We next examined whether the rescue of EC elongation was reflected by improved F-actin rearrangements. We treated *tln1*^*uq1al-/-*^mutants and siblings with Jasp for 5 hours prior to time-lapse imaging. As previously observed, junctional F-actin was static in 0.1% DMSO treated *tln1*^*uq1al-/-*^embryos (Fig. 6D, E and Movie 5), however F-actin displacements were more pronounced in Jasp treated *tln1*^*uq1al-/-*^embryos (Fig. 6D, F and Movie 6). Taken together, these experiments show that stabilising the F-actin network can result in sufficient acto-myosin activity that is required to elongate FA-deficient ECs and for junctions to linearise. Whether compensation occurs via immature cell-ECM adhesive structures or whether elongation is mainly driven by JBLs at cell-cell junctions remains to be determined.

**Figure 6:**
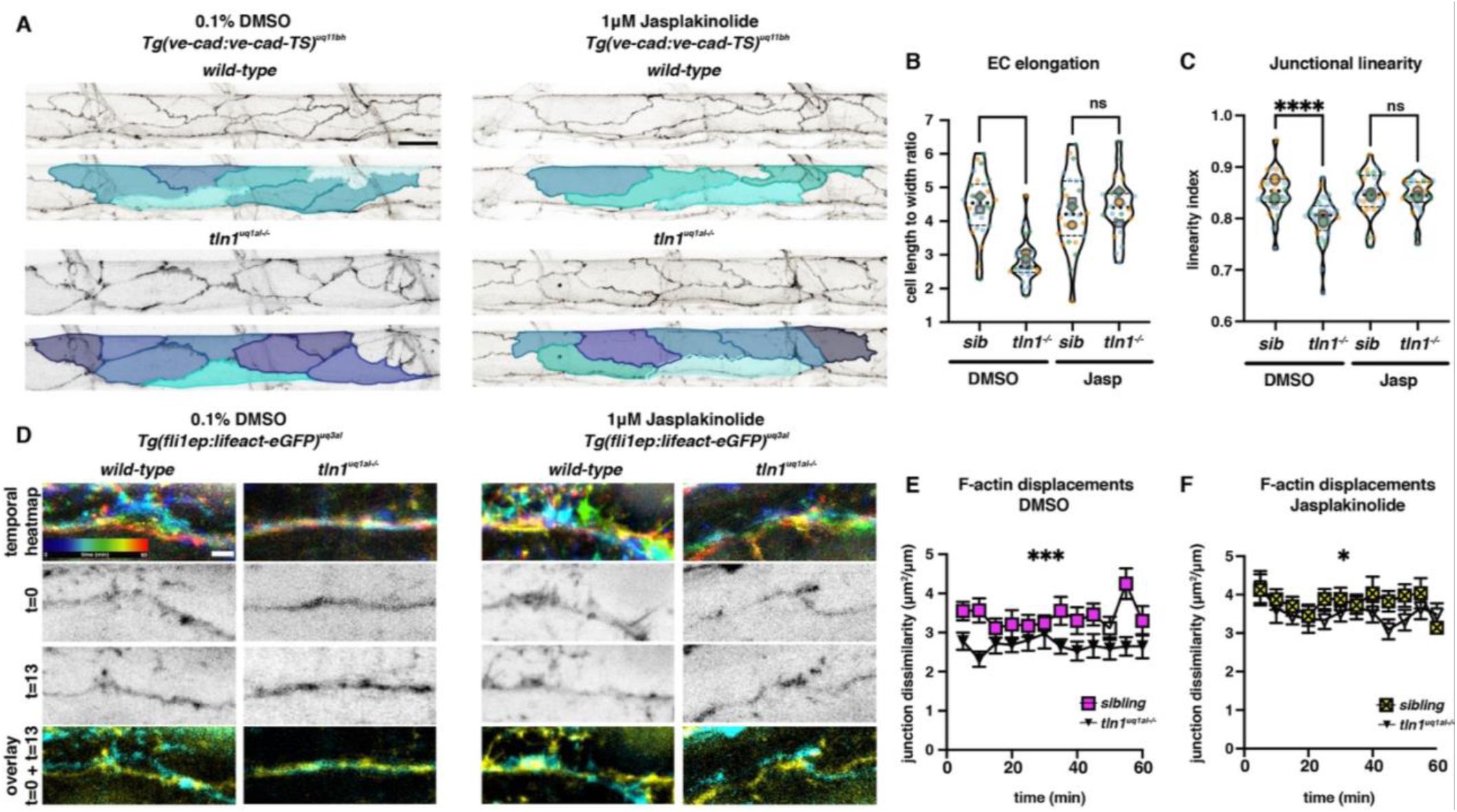
Enhancing F-actin stability can restore EC elongation, junctional linearity, and F-actin rearrangements in *tln1*^*uq1al-/-*^mutants. (A) Maximum projection of the dorsal aorta at 50 hpf, directly after treatment with either 0.1% DMSO (control, left) or Jasplakinolide (Jasp, right) [1μM], showing impaired EC elongation in DMSO treated *tln1*^*uq1al-/-*^mutants (bottom left) which is rescued when mutants have been exposed to Jasp (bottom right). The colours present a spectrum, lighter colours indicate more elongated ECs. Scale bar = 25µm. (B) Quantification of EC elongation at 50 hpf, showing a phenotypic rescue of EC elongation in Jasp treated *tln1*^*uq1al-/-*^mutants, n=3 biological replicates, n=27 siblings and n=25 *tln1*^*uq1al-/-*^mutants in DMSO treated group and n=27 siblings and n=22 *tln1*^*uq1al-/-*^mutants in Jasp treated group. (C) Quantification of cell-cell junction linearity at 50 hpf, showing significant rescue of linearisation in Jasp treated *tln1*^*uq1al-/-*^mutants, n=3 biological replicates, n=27 siblings and n=25 *tln1*^*uq1al-/-*^mutants in DMSO treated group and n=27 siblings and n=22 *tln1*^*uq1al-/-*^mutants in Jasp treated group. (D) Stills from time-lapse imaging (Movie 5 and 6) of cortical F-actin displacements in 0.1% DMSO (control, left) or Jasp [1μM] treated (right) wild-type siblings and and *tln1*^*uq1al-/-*^mutant embryos. Each column of images is organised as follows; Top: temporal colour-coded projection of 13 timepoints imaged during the hour time-lapse. Middle: Greyscale images of F-actin expression at the start (t=0) and the end of the movie (t=13). Bottom: overlay of t=0 (cyan) and t=13 (yellow) showing F-actin displacement. Scale bar = 5μm (E) Quantification of cortical F-actin displacements over the course of a one-hour time-lapse movie in 0.1% DMSO treated siblings (magenta) and *tln1*^*uq1al-/-*^mutants (black). Scale bar = 10µm (F) Quantification of cortical F-actin displacements over the course of a one-hour time-lapse movie in Jasp treated siblings (magenta) and *tln1*^*uq1al-/-*^mutants (black) showing a partial rescue when compared to E, n=3 biological replicates, n=12 siblings and n=12 *tln1*^*uq1al-/-*^mutants in DMSO treated group and n=14 siblings and n=10 *tln1*^*uq1al-/-*^mutants in Jasp treated group. In B and C replicate averages are depicted by large circles. Smaller circles present individual data points of each replicate (colour matched).

## DISCUSSION

This study reports the first live *in vivo* analysis of FA deficient endothelial cells during early vascular morphogenesis. By using a novel, FA-deficient, *talin1* zebrafish mutant model, we show that vasculogenesis and lumenisation of the DA and PCV occurs normally, unlike global and early EC-specific Talin1 knockout mice, where severe defects in in vasculogenesis and lumenisation have been observed^4,36^. Talin1 protein has previously been suggested to be extremely stable and functions sufficiently for a period of time in ECs that have undergone Talin1 deletion^4^. One explanation for the difference in severity of these phenotypes might be that Talin1 protein and/or mRNA is maternally deposited at sufficient levels in our zebrafish model to allow these early processes of vascular morphogenesis to occur^50^. The fact that the DA and PCV develop normally, combined with sufficient cardiac function, conveys a significant benefit, since this allowed us to investigate FA-deficient ECs in flow pressured vasculature. Using a novel endothelial restricted Vinculin transgenic line, we were able to verify a loss of FAs in *tln1*^*uq1al-/-*^mutants. Live imaging of Vinculin revealed extremely dynamic behaviour of FAs in wild-type ECs, which has not been shown *in vivo* previously. This line therefore will provide a unique resource to the cardio-vascular research community for future studies into FA function.

We uncovered that FA function is required to facilitate EC elongation and cell-cell junction linearisation in both angiogenic ISVs and in the main axial artery, the DA. This reduced capacity of ECs to elongate resulted in a failure of the vascular network to remodel and mature. The immature ISVs frequently presented as unicellular tubes. We hypothesise that such unicellular regions would be more fragile once exposed to blood flow, due to high pressure inflicted onto a single EC, instead of being distributed over multiple cells. In support of this, we found that a significant proportion of the ISVs detached from the main vascular network at 2 dpf. A further explanation for the failure of ECs to reorganise appropriately in ISVs might be through a weakening of cell-cell junctions via FA-cadherin crosstalk. VE-cadherin at cell-cell junctions has previously been shown to be essential for EC elongation during ISV multicellular tube formation^41,45,48^. Together this failure of ISVs to remodel can explain the severe systemic haemorrhaging that has been observed in mouse models where Talin1 was depleted specifically during the developmental time window of angiogenesis^4^.

We further showed that Talin1 was also necessary for EC elongation and cell-cell junction linearisation of larger calibre vessels, specifically in the DA, and that this role was endothelial cell-autonomous. Previous studies have demonstrated that EC elongation is an essential process that occurs during DA maturation^40,42^. When elongation is impaired, the DA will fail to narrow over time which leads to systemic perfusion defects^42^. Concurrently, the EC junctions also continued to display an immature irregular morphology, instead of linearising from 2 to 3 dpf^40,42^. This irregular VE-cadherin patterning is in agreement with previous observations made from EC specific integrin-β1 and Talin1 knockout mice^22,27^, where VE-cadherin was found to be distributed over a wider area at the cell-cell junctions. By using small molecule inhibitors that alter actin polymerisation, we showed that inhibition of F-actin polymerisation phenocopied *tln1*^*uq1al-/-*^mutants, with a significant reduction in EC elongation and junction linearisation. Furthermore, enhancement of actin polymerisation could restore EC elongation and junction linearisation in *tln1*^*uq1al-/-*^mutants. Together with our observation that Talin1 deficiency results in a loss of Vinculin clusters both at FAs and cell-cell junctions, we suggest that significant FA-cadherin crosstalk exists via acto-myosin interactions to control EC elongation and cell-cell junction linearisation.

Precisely where and when acto-myosin acts to allow EC elongation during vascular remodelling remains to be determined. Notably, these cellular processes might also be dependent on Talin1 function at cell-cell junctions, given that Pulous and colleagues have shown previously that Talins are expressed at cell-cell junctions in cultured Human umbilical cord venous ECs (HUVECs) and human dermal microvascular endothelial cells (HDMVECs)^22^. Whether Talin1 is also localised at endothelial cell-cell junctions *in vivo* however remains to be determined. Phenotypic rescue of EC elongation in *tln1*^*uq1al-/-*^by the actin stabiliser Jasplakinolide also support the idea that redundant FAs might exist, linked to actin via other adaptor proteins such as kindlins.

Overall, our findings reveal a previously unappreciated role for Talin1, and more broadly for FAs, in EC elongation and junction linearisation. The failure of these process resulted in unstable vasculature with irregular cell-cell junctions, explaining the occurrence of haemorrhaging vasculature across a range of FA deficient animal models.

## MATERIALS AND METHODS

### Zebrafish stocks and husbandry

All animal work adhered to the guidelines of the animal ethics committee at the University of Queensland. Published transgenic lines used for this work are *Tg(ve-cad:ve-cadTS)*^*uq11bh* 40^ and *Tg(kdrl:Hsa*.*HRAS-mCherry)*^*s916* 51^. For live imaging purposes, all embryos were treated with 0.0003% phenylthiourea (PTU) from 24 hpf to prevent pigmentation and anaesthetized in Tricaine (Sigma Aldrich E10521-50G, 0.08mg/ml).

### Cloning and Transgenesis

To establish *Tg(fli1ep:Vinculinb-eGFP)*^*uq2al*^ we amplified Vinculinb cDNA and cloned into the Gateway pME vector (pDON-221) using Gateway technology^52^. Oligos used to amplify Vinculinb: Vinculinb-Forward: 5′-**GGGGACAAGTTTGTACAAAAAAGCAGGCT** ACCATGCCGGTTTTCCACACG-3′ (gateway homology arm = bold, kozac sequence = underlined), and Vinculinb-Reverse: 5′-**GGGGACCACTTTGTACAAGAAAGCTGGGTA** CTGGTACCAGGGTGTCTTGC -3′ (gateway homology arm = bold). Subsequently a Gateway LR reaction was performed combining a p5E-fli1ep, pME-Vinculinb and p5E-eGFP placing the final *fli1ep:Vinculinb-eGFP* sequence into pDestTol2pA2AC (containing the α-crystallin promoter driving GFP in the zebrafish lens). *Tg*(*fli1ep:lifeact-eGFP)*^*uq3al*^ was generated by injection of a published plasmid^1^ (gift from Holger Gerhardt and Li-Kun Phng). For transgenesis we injected 1 nl of purified plasmid DNA (100 ng/μl) into single-cell stage embryos together with tol2 transposase mRNA (25 ng/μl). Embryos expressing Vinculin-eGFP or Lifeact-eGFP mosaically were selected and raised to adulthood. Adults were screened for germline transmission to generate stable transgenic lines.

### Genome editing and Genotyping

CRISPR genome editing for *talin1* was performed as previously described^53^. Oligo sequence used for guide RNA transcription: 5′-*taatacgactcactata*GGGCGGACGCTGCGGGAGCGTTTTAGAGCTAGAAATAGCAAG –3′ (T7 site for transcription = italics and guide RNA sequence = underlined)

To genotype the *tln1*^*uq1al-/-*^allele we amplified a 124bp region with primers flanking the mutation site. Amplified products were separated using 2% sodium borate agarose gels. Genotypes were distinguished as follows; heterozygotes: two bands of 124bp and 107bp, homozygous wild-type: single band of 124bp and homozygous mutant: single band of 107bp. Primer sequences for genotyping PCR: Talin1-Forward: 5’-CGCTGATTGGGATTTTAATGTGTATTCAG -3’, Talin1-Reverse: 5’-CTGGTCTGAGTAGAAGAACTTTCTCC - 3’

### Quantitative real-time polymerase chain reaction

RNA was extracted from 2 dpf whole embryos using the Directzol RNA mini kit (Zymo). cDNA was synthesised and amplified using Thermo Fisher SuperScript II Reverse Transcriptase Kit. qPCR was performed on Applied Biosystems Viia 7 384 well real-time PCR machine. There were 3.75μL 2x SYBR Green Master Mix, 3.25μL cDNA (1/20 dilution). Fold changes were calculated relative to housekeeping genes *rpl13* and *hprt*.

### Embryonic Transplantation

Transplantation was performed as previously described^51^. Donor embryos were derived from *tln1*^*uq1al+/-*^inter crosses (with transgenic background *Tg(ve-cad:ve-cadTS)*^*uq11bh*^). Cells were transferred at blastula stages into *Tg(kdrl:Hsa*.*HRAS-mCherry)*^*s916*^ wild-type hosts using CellTram 4R Oil hydrolic microinjector (Eppendorf). Donor embryos were genotyped immediately after transplantation.

### Chemical treatments

For CK666 (Arp2/3 inhibitor) treatments, embryos were incubated in E3 medium containing CK666 [75μM] in 0.4% DMSO or 0.4% DMSO only as a negative control from 24 hpf till 50 hpf. For Jasplakinolide (Jasp) embryos were incubated in E3 medium containing Jasp [1μM] in 0.1% DMSO or 0.1% DMSO in E3 as a negative control 43 hpf until 48 hpf and subsequently imaged from 48 till 49 hpf.

### Image acquisition

Embryos were mounted in 35mm glass bottom dishes using 0.7% low melting point agarose. Confocal z-stacks were acquired on a LSM 710 Meta BiG (Zeiss) and a LSM 880 FastAiry (Zeiss) using a LD C-Apochromat W/ 40x 1.1 NA objective. Timelapse imaging was performed on a Dragonfly Spinning Disc microscope (Andor) using Apo λ LWD W/ 40x 1.15 NA. Brightfield images of the whole-mount zebrafish embryos were taken on a Nikon SMZ1270i. Movie capturing real-time blood flow (Movie 2) was captured on a Nikon Ti-E inverted microscope using a Plan Apochromat λ/40x 0.95 NA objective.

### Image processing and quantifications

All image processing and analysis was performed by using the Fiji, ImageJ (National Institutes of Health)^54^. To ensure consistency, all quantifications of ECs in the ISVs and the DA were performed in a specific section of the trunk, above the yolk extension.

#### Vinculinb positive clusters

Median filter of 10 pixels radius was applied to the images and the images were thresholded to select for Vinculinb expression. Analyse particles (Fiji, ImageJ) was applied to create a masks for junctional and cytoplasmic Vinculinb-clusters (cytoplasmic clusters: <5μm^2^, junctional clusters: >5μm^2^). The masks were applied to the original images and the “Find maxima” function (prominence >15) was used to count the number of clusters in each area.

#### Cell-cell junction width

Median filter of 10 pixels radius was applied to the images and the images were thresholded to select for Vinculinb expression. Analyse particles was performed on the thresholded images to create mask for junctional area based on particle size (>5μm^2^). The original image was masked and thresholded to select for vinculin expression. The threshold area was measured as the junction area. The threshold images were then skeletonized, and the length of the skeleton was measured as the junction length. The junction area was divided by the junction length to yield the junction width. Filopodia number and length: The number of filopodia within the first 10μm of the ISV tip was counted manually. The length of filopodia within the first 10μm of the ISV sprout was measured using the freehand line tool (Fiji, ImageJ).

#### Number of DLAV plexus branches

The number of dorsal vessel branches at the DLAV plexus within the last two somites above the yolk extension was quantified manually.

#### Number of disconnected ISVs

Number of disconnected ISVs within the last three ISVs above the yolk extension was counted manually in each embryo.

#### ISV sprout length

ISV sprout length was measured by using the line tool (Fiji, ImageJ) and drawing a line from the base of the sprout at the level of the DA to the tip of the sprout. Percentage of multicellular ISV: We employed previously described methodology to distinguish unicellular from multicellular sections of the ISVs^45^. The percentage was calculated by the number of multicellular ISVs out of the three ISVs captured in each image.

#### Endothelial cell elongation

Using the freehand tool (Fiji, ImageJ), a region of interest (ROI) was drawn to outline individual ECs (based on *Tg(ve-cad:ve-cadTS)*^*uq11bh*^ expression). We subsequently measured cell elongation by using the “fit ellipse” shape measurement function (Fiji, ImageJ). Cellular ellipticity was calculated by dividing the ellipse length by the ellipse width.

#### Junctional linearity

A junctional linearity index was calculated as previously described^40^

#### F-actin displacement

We measured the area by which F-actin expression had shifted between two consecutive time-points of the movies. To do this, we generated an overlay of *Tg*(*fli1ep:lifeact-eGFP)*^*uq3al*^ expression and measured the surface area between the two. We subsequently normalised this by the average junction length of the two time points.

### Statistical analysis

We performed all statistical analysis using Prism 9 (GraphPad). Prism Violin plots were combined with “all dot plots” using the web based software SuperPlotsOfData^55^, providing a transparent display and quantitative comparison between experimental replicates. ROUT method was applied to test normal distribution of the data points. When the data were normally distributed a Student’s t-test was used for comparison of two means. When the data did not follow a normal distribution, the Mann–Whitney test was used for comparison of two means. The threshold for significance was taken as p < 0.05 and all the data are represented as mean ± s.e.m For each experimental comparison, embryos were randomly distributed, and sibling matched.

## ACKNOWLEDGEMENTS

We thank Dr. S. Stehbens and Dr. L-K. Phng for critical reading of this manuscript. We thank Prof. B. Collins for assistance with Alphafold protein folding predictions. We thank Prof. H. Gerhardt and Dr. L-K. Phng for kindly providing the DNA construct that we used to generate the *Tg(fli1ep:lifeact-eGFP)*^*uq3al*^ strain. Imaging was performed in the Australian Cancer Research Foundation’s Dynamic Imaging Facility at IMB (established with the generous support of the ACRF). We thank the UQBR aquatics facility staff for animal caretaking.

## COMPETING INTERESTS

The authors declare no competing financial interests

## FUNDING

T.C.Y.C. was supported by a UQ RS2 postdoctoral fellowship, A.S.Y by a NHMRC Senior Principal Research Fellowship (APP1136592) and B.M.H. by a NHMRC Research Fellowship (1155221). This research was supported by an ARC Discovery Project grant (DP200100737) and a NHMRC Project Grant (2002436).

## DATA AVAILABILITY

All the relevant data supporting the findings are available from the corresponding author on reasonable request.

## Notes

### Competing Interest Statement

The authors have declared no competing interest.

## REFERENCES

1 Phng, L. K., Stanchi, F. & Gerhardt, H. Filopodia are dispensable for endothelial tip cell guidance. Development 140, 4031–4040, doi:10.1242/dev.097352 (2013).

2 Campinho, P., Vilfan, A. & Vermot, J. Blood Flow Forces in Shaping the Vascular System: A Focus on Endothelial Cell Behavior. Front Physiol 11, 552, doi:10.3389/fphys.2020.00552 (2020).

3 Fonseca, C. G., Barbacena, P. & Franco, C. A. Endothelial cells on the move: dynamics in vascular morphogenesis and disease. Vasc Biol 2, H29–H43, doi:10.1530/VB-20-0007 (2020).

4 Monkley, S. J. et al. Endothelial cell talin1 is essential for embryonic angiogenesis. Dev Biol 349, 494–502, doi:10.1016/j.ydbio.2010.11.010 (2011).

5 Carlson, T. R., Hu, H., Braren, R., Kim, Y. H. & Wang, R. A. Cell-autonomous requirement for beta1 integrin in endothelial cell adhesion, migration and survival during angiogenesis in mice. Development 135, 2193–2202, doi:10.1242/dev.016378 (2008).

6 Lei, L. et al. Endothelial expression of beta1 integrin is required for embryonic vascular patterning and postnatal vascular remodeling. Mol Cell Biol 28, 794–802, doi:10.1128/MCB.00443-07 (2008).

7 Tanjore, H., Zeisberg, E. M., Gerami-Naini, B. & Kalluri, R. Beta1 integrin expression on endothelial cells is required for angiogenesis but not for vasculogenesis. Dev Dyn 237, 75–82, doi:10.1002/dvdy.21385 (2008).

8 Zovein, A. C. et al. Beta1 integrin establishes endothelial cell polarity and arteriolar lumen formation via a Par3-dependent mechanism. Dev Cell 18, 39–51, doi:10.1016/j.devcel.2009.12.006 (2010).

9 Kechagia, J. Z., Ivaska, J. & Roca-Cusachs, P. Integrins as biomechanical sensors of the microenvironment. Nat Rev Mol Cell Biol 20, 457–473, doi:10.1038/s41580-019-0134-2 (2019).

10 Sun, Z., Guo, S. S. & Fassler, R. Integrin-mediated mechanotransduction. J Cell Biol 215, 445–456, doi:10.1083/jcb.201609037 (2016).

11 Calderwood, D. A., Campbell, I. D. & Critchley, D. R. Talins and kindlins: partners in integrin-mediated adhesion. Nat Rev Mol Cell Biol 14, 503–517, doi:10.1038/nrm3624 (2013).

12 Goult, B. T., Yan, J. & Schwartz, M. A. Talin as a mechanosensitive signaling hub. J Cell Biol 217, 3776–3784, doi:10.1083/jcb.201808061 (2018).

13 Klapholz, B. & Brown, N. H. Talin - the master of integrin adhesions. J Cell Sci 130, 2435–2446, doi:10.1242/jcs.190991 (2017).

14 Harburger, D. S. & Calderwood, D. A. Integrin signalling at a glance. J Cell Sci 122, 159–163, doi:10.1242/jcs.018093 (2009).

15 Goult, B. T. et al. Structure of a double ubiquitin-like domain in the talin head: a role in integrin activation. EMBO J 29, 1069–1080, doi:10.1038/emboj.2010.4 (2010).

16 Rupp, P. A. & Little, C. D. Integrins in vascular development. Circ Res 89, 566–572, doi:10.1161/hh1901.097747 (2001).

17 Dejana, E., Lampugnani, M. G., Giorgi, M., Gaboli, M. & Marchisio, P. C. Fibrinogen induces endothelial cell adhesion and spreading via the release of endogenous matrix proteins and the recruitment of more than one integrin receptor. Blood 75, 1509–1517, doi:10.1182/blood.V75.7.1509.bloodjournal7571509 (1990).

18 Wu, M. H., Ustinova, E. & Granger, H. J. Integrin binding to fibronectin and vitronectin maintains the barrier function of isolated porcine coronary venules. Journal of Physiology 532, 785–791, doi:10.1111/j.1469-7793.2001.0785e.x (2001).

19 Malinin, N. L., Pluskota, E. & Byzova, T. V. Integrin signaling in vascular function. Curr Opin Hematol 19, 206–211, doi:10.1097/MOH.0b013e3283523df0 (2012).

20 van der Flier, A. et al. Endothelial alpha5 and alphav integrins cooperate in remodeling of the vasculature during development. Development 137, 2439–2449, doi:10.1242/dev.049551 (2010).

21 Kopp, P. M. et al. Studies on the morphology and spreading of human endothelial cells define key inter-and intramolecular interactions for talin1. Eur J Cell Biol 89, 661–673, doi:10.1016/j.ejcb.2010.05.003 (2010).

22 Pulous, F. E. et al. Talin-dependent integrin activation is required for endothelial proliferation and postnatal angiogenesis. Angiogenesis 24, 177–190, doi:10.1007/s10456-020-09756-4 (2021).

23 Wu, Q. et al. Talin1 is required for cardiac Z‐disk stabilization and endothelial integrity in zebrafish. FASEB Journal 29, 4989–5005, doi:10.1096/fj.15-273409 (2015).

24 Mui, K. L., Chen, C. S. & Assoian, R. K. The mechanical regulation of integrin-cadherin crosstalk organizes cells, signaling and forces. J Cell Sci 129, 1093–1100, doi:10.1242/jcs.183699 (2016).

25 Schwartz, M. A. & DeSimone, D. W. Cell adhesion receptors in mechanotransduction. Curr Opin Cell Biol 20, 551–556, doi:10.1016/j.ceb.2008.05.005 (2008).

26 Weber, G. F., Bjerke, M. A. & DeSimone, D. W. Integrins and cadherins join forces to form adhesive networks. J Cell Sci 124, 1183–1193, doi:10.1242/jcs.064618 (2011).

27 Yamamoto, H. et al. Integrin β1 controls VE-cadherin localization and blood vessel stability. Nature Communications 6, doi:10.1038/ncomms7429 (2015).

28 Chen, K. D. et al. Mechanotransduction in response to shear stress. Roles of receptor tyrosine kinases, integrins, and Shc. J Biol Chem 274, 18393–18400, doi:10.1074/jbc.274.26.18393 (1999).

29 Orr, A. W., Ginsberg, M. H., Shattil, S. J., Deckmyn, H. & Schwartz, M. A. Matrix-specific suppression of integrin activation in shear stress signaling. Mol Biol Cell 17, 4686–4697, doi:10.1091/mbc.e06-04-0289 (2006).

30 Tzima, E., del Pozo, M. A., Shattil, S. J., Chien, S. & Schwartz, M. A. Activation of integrins in endothelial cells by fluid shear stress mediates Rho-dependent cytoskeletal alignment. EMBO J 20, 4639–4647, doi:10.1093/emboj/20.17.4639 (2001).

31 Tzima, E. et al. A mechanosensory complex that mediates the endothelial cell response to fluid shear stress. Nature 437, 426–431, doi:10.1038/nature03952 (2005).

32 Jumper, J. et al. Highly accurate protein structure prediction with AlphaFold. Nature 596, 583–589, doi:10.1038/s41586-021-03819-2 (2021).

33 Tunyasuvunakool, K. et al. Highly accurate protein structure prediction for the human proteome. Nature 596, 590–596, doi:10.1038/s41586-021-03828-1 (2021).

34 El-Brolosy, M. A. et al. Genetic compensation triggered by mutant mRNA degradation. Nature 568, 193–197, doi:10.1038/s41586-019-1064-z (2019).

35 Yamamoto, H. et al. Integrin beta1 controls VE-cadherin localization and blood vessel stability. Nat Commun 6, 6429, doi:10.1038/ncomms7429 (2015).

36 Pulous, F. E., Grimsley-Myers, C. M., Kansal, S., Kowalczyk, A. P. & Petrich, B. G. Talin-Dependent Integrin Activation Regulates VE-Cadherin Localization and Endothelial Cell Barrier Function. Circ Res 124, 891–903, doi:10.1161/CIRCRESAHA.118.314560 (2019).

37 Wu, Q. et al. Talin1 is required for cardiac Z-disk stabilization and endothelial integrity in zebrafish. FASEB J 29, 4989–5005, doi:10.1096/fj.15-273409 (2015).

38 Iida, A., Wang, Z., Hirata, H. & Sehara‐Fujisawa, A. Integrin β1 activity is required for cardiovascular formation in zebrafish. Genes to Cells 23, 938–951, doi:10.1111/gtc.12641 (2018).

39 Han, M. K. L., van der Krogt, G. N. M. & de Rooij, J. Zygotic vinculin is not essential for embryonic development in zebrafish. PLoS One 12, e0182278, doi:10.1371/journal.pone.0182278 (2017).

40 Lagendijk, A. K. et al. Live imaging molecular changes in junctional tension upon VE-cadherin in zebrafish. Nat Commun 8, 1402, doi:10.1038/s41467-017-01325-6 (2017).

41 Phng, L. K. & Belting, H. G. Endothelial cell mechanics and blood flow forces in vascular morphogenesis. Semin Cell Dev Biol, doi:10.1016/j.semcdb.2021.06.005 (2021).

42 Sugden, W. W. et al. Endoglin controls blood vessel diameter through endothelial cell shape changes in response to haemodynamic cues. Nat Cell Biol 19, 653–665, doi:10.1038/ncb3528 (2017).

43 Bentley, K. et al. The role of differential VE-cadherin dynamics in cell rearrangement during angiogenesis. Nat Cell Biol 16, 309–321, doi:10.1038/ncb2926 (2014).

44 Perryn, E. D., Czirok, A. & Little, C. D. Vascular sprout formation entails tissue deformations and VE-cadherin-dependent cell-autonomous motility. Dev Biol 313, 545–555, doi:10.1016/j.ydbio.2007.10.036 (2008).

45 Sauteur, L. et al. Cdh5/VE-cadherin promotes endothelial cell interface elongation via cortical actin polymerization during angiogenic sprouting. Cell Rep 9, 504–513, doi:10.1016/j.celrep.2014.09.024 (2014).

46 Cao, J. et al. Polarized actin and VE-cadherin dynamics regulate junctional remodelling and cell migration during sprouting angiogenesis. Nat Commun 8, 2210, doi:10.1038/s41467-017-02373-8 (2017).

47 Phng, L. K. et al. Formin-mediated actin polymerization at endothelial junctions is required for vessel lumen formation and stabilization. Dev Cell 32, 123–132, doi:10.1016/j.devcel.2014.11.017 (2015).

48 Paatero, I. et al. Junction-based lamellipodia drive endothelial cell rearrangements in vivo via a VE-cadherin-F-actin based oscillatory cell-cell interaction. Nat Commun 9, 3545, doi:10.1038/s41467-018-05851-9 (2018).

49 Holzinger, A. Jasplakinolide: an actin-specific reagent that promotes actin polymerization. Methods Mol Biol 586, 71–87, doi:10.1007/978-1-60761-376-3_4 (2009).

50 Harvey, S. A. et al. Identification of the zebrafish maternal and paternal transcriptomes. Development 140, 2703–2710, doi:10.1242/dev.095091 (2013).

51 Hogan, B. M. et al. Ccbe1 is required for embryonic lymphangiogenesis and venous sprouting. Nat Genet 41, 396–398, doi:10.1038/ng.321 (2009).

52 Hartley, J. L., Temple, G. F. & Brasch, M. A. DNA cloning using in vitro site-specific recombination. Genome Res 10, 1788–1795, doi:10.1101/gr.143000 (2000).

53 Gagnon, J. A. et al. Efficient mutagenesis by Cas9 protein-mediated oligonucleotide insertion and large-scale assessment of single-guide RNAs. PLoS One 9, e98186, doi:10.1371/journal.pone.0098186 (2014).

54 Schindelin, J. et al. Fiji: an open-source platform for biological-image analysis. Nature Methods 9, 676, doi:10.1038/nmeth.2019 (2012).

55 Lord, S. J., Velle, K. B., Mullins, R. D. & Fritz-Laylin, L. K. SuperPlots: Communicating reproducibility and variability in cell biology. J Cell Biol 219, doi:10.1083/jcb.202001064 (2020).

