## Supplementary Figures for "Dynamically regulated Focal adhesions coordinate endothelial cell remodelling in developing vasculature"

### Supplementary Figure 1

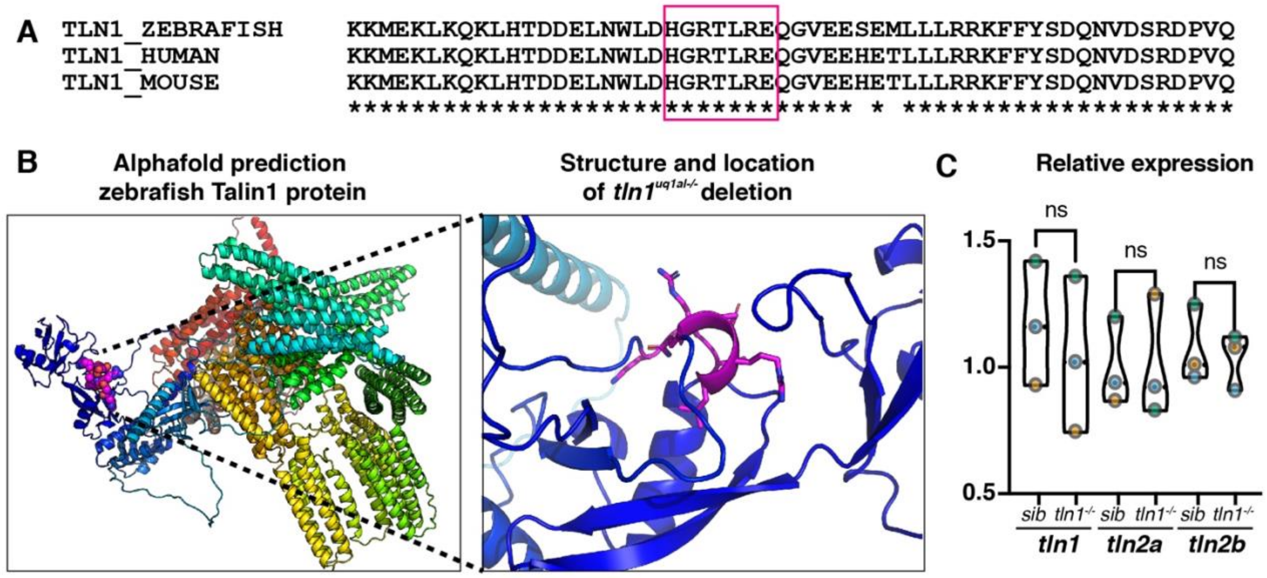


**Supplementary Figure 1: Sequence and structure of deleted amino acids in *tln1*$^{uq1al-/-}$ mutants**

(A) Multi-species protein alignment of the CRISPR-Cas9 targeted region within the highly conserved F1 FERM domain of Talin1 in zebrafish (top), human (middle) and mouse (bottom). The box indicates the amino acids that have been deleted in our *tln1*$^{uq1al-/-}$ mutant line. (B) Left: Schematic model showing the predicted structure of zebrafish Talin1 based on AlphaFold entry A0A0R4IDZ8. The model is coloured in a spectrum from blue to red from N- to C-terminus. Blue presents the N-terminal FERM domain and in magenta are the amino acids deleted in *tln1*$^{uq1al-/-}$ mutants. Right: Detailed view showing the wild-type structure and orientation of amino acids (magenta) that are not present in our *tln1*$^{uq1al-/-}$ mutants. (C) Quantitative RT-PCR analysis of *tln1*, *tln2a* and *tln2b* expression levels in sibling and *tln1*$^{uq1al-/-}$ mutant embryos at 2 dpf, n=3 biological replicates, n=30 siblings and n=30 *tln1*$^{uq1al-/-}$ mutants. Replicate averages are depicted by the circles.

#### Supplementary Figure 2

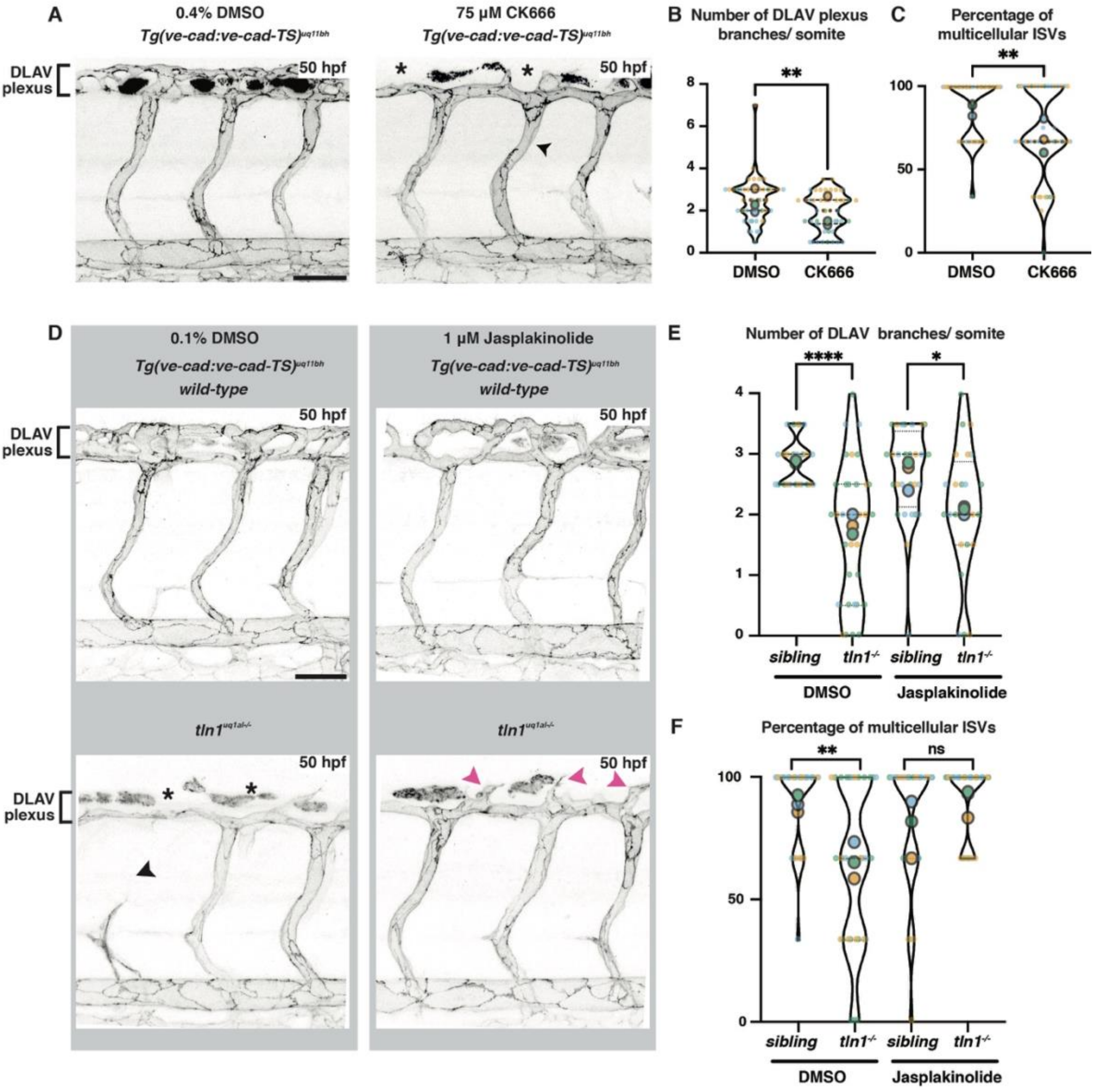


**Supplementary Figure 2: Vessel network abnormalities in CK666 treated animals and improved vessel morphogenesis in Jasplakinolide treated *tln1*$^{uq1al-/-}$ mutants**

(A) CK666 treated wild-type embryos develop phenotypes that are similar to *tln1*$^{uq1al-/-}$ mutants, including compromised DLAV plexus formation (asterisk) and the more frequent occurrence of immature unicellular tubes (arrowhead). Scale bar = 50 µm (B) Quantification of the number of DLAV branches per somite, n=3 biological replicates, n=48 DMSO and n=48 CK666 treated embryos. (C) Quantification of the percentage of multicellular ISVs per 3 somites, n=3 biological replicates, n=48 DMSO and n=48 CK666 treated embryos. (D) Maximum projection of the trunk vasculature at 50 hpf, directly after treatment with either DMSO (control, left) or Jasp (right), showing more DLAV plexus branches in Jasp *tln1*$^{uq1al-/-}$ mutants (asterisk versus magenta arrowheads). Scale bar = 50 µm (B) Quantification of the number of DLAV branches per somite in DMSO versus Jasp treated embryos, n=3 biological replicates, n=25 siblings and n=33 *tln1*$^{uq1al-/-}$ mutants in the DMSO treated group and n=28 siblings and n=24 *tln1*$^{uq1al-/-}$ mutants in the Jasp treated group. (C) Quantification of the percentage of multicellular ISVs per 3 somites in DMSO versus Jasp treated embryos, n=3 biological replicates, n=25 siblings and n=33 *tln1*$^{uq1al-/-}$ mutants in the DMSO treated group and n=28 siblings and n=24 *tln1*$^{uq1al-/-}$ mutants in the Jasp treated group. In all graphs replicate averages are depicted by large circles. Smaller circles present individual data points of each replicate (colour matched).

**MOVIE CAPTIONS**

**Supplementary Movie 1: Time lapse movie showing endothelial Vinculin dynamics *in vivo***

**Top:** Dorsal aorta (DA) of a wild-type embryo, expressing *Tg(fli1ep:Vinculinb-eGFP)*$^{uq2al}$, showing dynamic Vinculin expression Focal Adhesions and cell-cell junctions over the course of the movie, from 50 hpf to 51 hpf.

**Bottom:** DA of a *tln1*$^{uq1al-/-}$ mutants, expressing *Tg(fli1ep:Vinculinb-eGFP)*$^{uq2al}$, showing a dramatic reduction of Focal Adhesions and diffuse cell-cell junctions expression, from 50 hpf to 51 hpf. Scale bar = 10μm.

**Supplementary Movie 2: Blood circulation in trunk vasculature**

Live recording of blood flow in the dorsal aorta (DA) of a wild-type embryo (top) and a *tln1*$^{uq1al-/-}$ mutant (bottom) at 2 dpf. Scale bar = 25μm.

**Supplementary Movie 3: Time-lapse imaging during ISVs remodelling**

Time-lapse imaging of ISVs in a wild-type embryo (left) and a *tln1*$^{uq1al-/-}$ mutant (right), showing disconnection of the ISV in the mutant. The endothelial F-actin marker line, *Tg(fli1ep:lifeact-eGFP)*$^{uq3al}$, was used to visualise the vasculature. Arrow indicates ECs disconnecting in the ISV of *tln1*$^{uq1al-/-}$ mutant. Movie taken from 30 hpf to 48 hpf. Scale bar = 25μm.

**Supplementary Movie 4: Impaired F-actin rearrangements in *tln1*$^{uq1al-/-}$ mutants**

Time-lapse imaging of cortical F-actin, visualised by *Tg(fli1ep:lifeact-eGFP)*$^{uq3al}$, near a cell-cell junction in a wild-type sibling (left) and a *tln1*$^{uq1al-/-}$ mutant (right). Movie was taken from 48 hpf to 49 hpf. Scale bar = 5μm.

**Supplementary Movie 5: 0.1% DMSO treated control**

Time-lapse imaging of cortical F-actin, visualised by *Tg(fli1ep:lifeact-eGFP)*$^{uq3al}$, near a cell-cell junction of a 0.1% DMSO treated wild-type sibling (left) and *tln1*$^{uq1al-/-}$ mutant (right). Embryos were treated for 5 hours prior to imaging. Movie was taken from 48 hpf to 49 hpf. Scale bar = 5μm.

**Supplementary Movie 6: Jasplakinolide induced rescue of F-acting rearrangements in *tln1*$^{uq1al-/-}$ mutants**

Time-lapse imaging of cortical F-actin, visualised by *Tg(fli1ep:lifeact-eGFP)*$^{uq3al}$, near a cell-cell junction of a Jasp [1μM] treated wild-type sibling (left) and *tln1*$^{uq1al-/-}$ mutant (right). Embryos were treated for 5 hours prior to imaging. Movie was taken from 48 hpf to 49 hpf. Scale bar = 5μm.
